# Modeling CRISPR-Cas13d on-target and off-target effects using machine learning approaches

**DOI:** 10.1101/2021.09.02.458773

**Authors:** Xiaolong Cheng, Zexu Li, Ruocheng Shan, Zihan Li, Lumen Chao, Jian Peng, Teng Fei, Wei Li

## Abstract

A major challenge in the application of the CRISPR-Cas13d (RfxCas13d, or CasRx) RNA editing system is to accurately predict its guide RNA (gRNA) dependent on-target and off-target effect. Here, we performed CRISPR-Cas13d proliferation screens that target protein-coding genes and long non-coding RNAs (lncRNAs), followed by a systematic modeling of Cas13d on-target efficiency and off-target viability effect. We first designed a deep learning model, named DeepCas13, to predict the on-target activity of a gRNA with high accuracy from its sequence and secondary structure. DeepCas13 outperforms existing methods and accurately predicts the efficiency of guides targeting both protein-coding and non-coding RNAs (e.g., circRNAs and lncRNAs). Next, we systematically studied guides targeting non-essential genes, and found that the off-target viability effect, defined as the unintended effect of guides on cell viability, is closely related to their on-target RNA cleavage efficiency. This finding suggests that these gRNAs should be used as negative controls in proliferation screens to reduce false positives, possibly coming from the unwanted off-target viability effect of efficient guides. Finally, we applied these models to our screens that included guides targeting 234 lncRNAs, and identified lncRNAs that affect cell viability and proliferation in multiple cell lines. DeepCas13 is freely accessible via http://deepcas13.weililab.org.

## Introduction

CRISPR-Cas13, including Cas13a, Cas13b, Cas13c, Cas13d (RfxCas13d, or CasRx) and the newly discovered Cas13X/Y (*1*), belongs to the type VI CRISPR-Cas system that exclusively target single-stranded RNA (ssRNA) (*2-4*). All Cas13 nucleases contain two HEPN domains as RNase to cleave RNAs or to process precursor crRNAs into mature crRNAs. Once activated by the single guide RNAs (sgRNAs) bearing complementarity sequences to the target RNA, Cas13 will cleave the target RNA and also nearby RNA molecules. Cas13 nucleases have also been engineered into efficient, multiplexable, and specific tools for the knockdown, editing and recognition of RNAs (and methylated RNAs like m^6^A RNA) in mammalian cells (*5-7*). In addition, Cas13 has been a central component of several rapid and sensitive methods for detecting viral infections including SARS-COV-2 (*8-11*).

A major challenge in the application of the CRISPR-Cas system (including Cas13) is to design single guide RNAs (sgRNAs) with high on-target efficiency and specificity. On the one hand, an accurate prediction of sgRNA efficiency would facilitate the optimized design of sgRNAs with maximized on-target efficiency (*i*.*e*., high sensitivity). On the other hand, understanding the specificity of Cas nucleases will help avoid the potential off-target effects, possibly due to the off-target cleavage at the DNA (for Cas9) or RNA (for Cas13) level, respectively, or due to its unwanted collateral cleavage to nearby mRNA molecules (Cas13) in many applications. For this reason, CRISPR screening has been a cost-effective approach to systematically investigate the efficiency and specificity of CRISPR-Cas9/Cas13, by examining the behaviors of a large number of guides in one single experiment. Guides in screens, including proliferation/viability screens and FACS-sorting screens, have been used to investigate factors that affect knockout efficiency (*12*), design guides that maximize activity and minimize off-target effects (*13*), and to train machine learning (*14*) and deep learning models (*15-17*) for a precise prediction of guide behaviors. Similar with Cas9 screens, Cas13d screens has been used to study the specificity and efficiency of Cas13d system (*18*), by examining a large number of tiling guides that span the gene of interest. Unlike CRISPR-Cas9 on-target activity prediction tools (*17, 19-21*) that only extract the spatial features of the sequence, models for Cas13 need to consider the secondary RNA structures of guides, which is a major factor for knockdown efficiency (*18*). However, there are some limitations to existing methods built for Cas13d efficiency prediction. First, the training dataset is based on FACS-sorting screens that measured the expression level of few specific genes, and it is unclear whether the corresponding model applies to guides targeting other genes and measuring other phenotypes (*e*.*g*., cell proliferation). Second, it is unclear whether such model, trained on guides targeting protein-coding genes, works on non-coding RNAs. Third, the off-target effect of Cas13d, mostly due to its collateral cleavage to mRNAs within the cell, is not fully explored.

Understanding the off-target effect of gene editing tools (*e*.*g*., TALEN, RNAi, Cas9, base editor), either dependent or independent from the guide, has been an important and challenging task in genome engineering. On the one hand, many assays have been developed to investigate the off-target editing outcomes of Cas9 guides (*22-26*), although most of these techniques are limited to report the effect of one sgRNA and none of them can be directly applied to Cas13 system. On the other hand, the non-specific toxicity effect has been examined on almost every gene editing tool. For example, the overexpression of short hairpin RNA (shRNA) damages cell viability by disrupting the miRNA processing in host cells (*27*). DNA double-strand breaks induced by Cas9 may also trigger DNA damage response (and subsequent cell death), particularly when guides target the amplified regions in the genome that results in stronger DNA damage (*28-30*). By examining thousands of Cas9 sgRNAs targeting non-functional, non-genic regions in the genome, the non-specific toxicity of Cas9 can be identified and mitigated in a screening manner (*31, 32*). The collateral RNA cleavage has been one of the major sources for Cas13 off-target effect (*5-7*), although such effect, especially on cell viability, has not been investigated systematically.

Here we systematically model the on-target efficiency and off-target viability effect of Cas13d (CasRx), using machine learning and deep learning approaches that are trained on large-scale Cas13d screening datasets. We first conducted CRISPR-Cas13d screens that contains 10,830 guides targeting essential/non-essential genes and long non-coding RNAs (lncRNAs). Combining this dataset with published studies, we obtained data from 22,599 Cas13d sgRNAs to systematically investigate the efficiency and specificity of Cas13d. We next designed DeepCas13, a deep learning-based model for predicting CRISPR-Cas13d on-target activity. DeepCas13 takes advantage of convolutional neural network and recurrent neural network to learn spatial-temporal features from the sequences and secondary structures of sgRNAs. DeepCas13 outperforms traditional machine learning methods and previous published tools, and demonstrates good performance in predicting guides targeting non-coding RNAs (*e*.*g*., circular RNAs and long non-coding RNAs). In addition, we systematically evaluated the off-target viability effect of Cas13d, by investigating guides targeting non-essential genes in the viability screens. We found that features determining the off-target viability effect of a guide are very similar with to features associated with on-target efficiency. Such effect can be mitigated in the Cas13d screens, using guides that target non-essential genes as negative controls. Compared with using non-targeting guides as negative controls, this approach greatly reduces false positives in the screen, a finding that resembles the use of non-essential guides in CRISPR/Cas9 screens. Finally, we applied these on-target and off-target models to CRISPR-Cas13d viability screens that target 234 lncRNAs, and identified putative lncRNAs whose perturbation reduces cell fitness in different cell lines. DeepCas13 is freely accessible via a web server at http://deepcas13.weililab.org/.

## Results

### Cas13d proliferation screen and a convolutional recurrent neural network for efficiency prediction

To systematically investigate the efficiency and specificity of Cas13d, we conducted a two-vector CRISPR/Cas13d proliferation screening experiment (Figure 1A; see Methods for details). The screening library contains 10,830 sgRNAs targeting 192 protein-coding genes and 234 lncRNAs, and the screening experiment was performed using a melanoma cell line A375. The library targets 94 known essential genes and 14 non-essential genes, which are identified from previous RNA interference and CRISPR screens (*33, 34*) and are confirmed to be essential (or non-essential) in A375, respectively (Supplementary Figure 1A), providing a unique dataset (3,934 guides in total and about 30 guides per gene) to model Cas13d sgRNA efficiencies. In addition, this dataset may potentially overcome the biases of previous tiling screening datasets, where guides that only target 2-3 genes are used. At the end of the screening, the abundance of Cas13d sgRNAs was evaluated using high-throughput sequencing, and the data analysis was performed using the MAGeCK algorithm we previously developed (*35*). Overall, the quality of the screen is high based on multiple quality control (QC) measurements (Supplementary Table S1), including sequencing depth (around 5.6 million reads per sample), the average number of reads per guide (over 300), the number of missing guides (less than 4), and the low Gini index values (less than 0.06) indicating the non-biased distribution of guides across different conditions. In addition, 20 out of 94 known essential genes are significantly depleted (FDR<10%; Supplementary Figure 1B, Supplementary File 1). These genes include MTOR (in mTOR pathway), TUBA1B (component of Tubulin complex), RPL4/RPS8 (ribosomal subunit), all known to be involved in essential functionalities of cellular functions. The guides targeting essential genes are strongly depleted, as expected, demonstrating the success of the screens (Supplementary Figure 1C).

**Figure 1.**
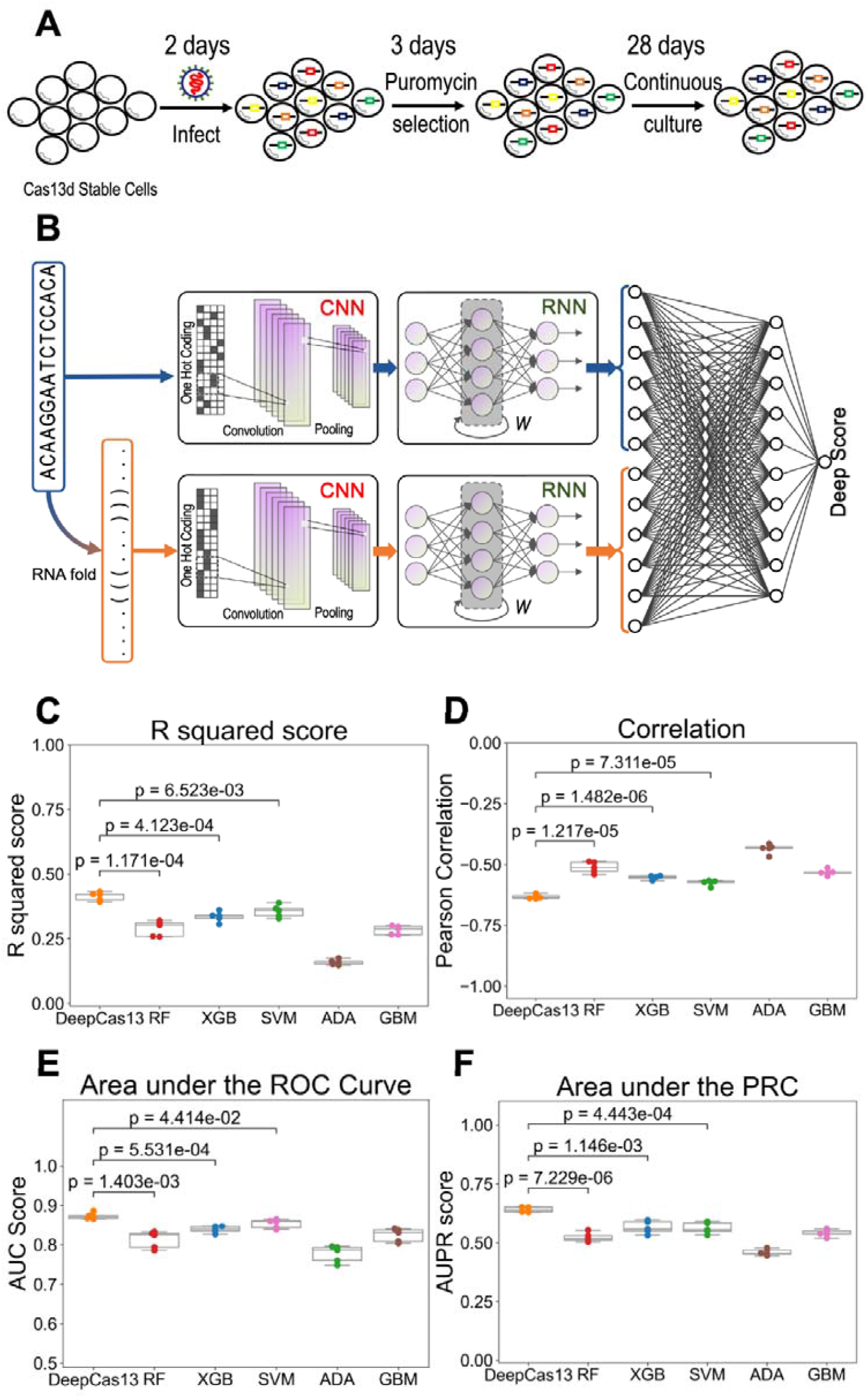
DeepCas13 predicts Cas13d on-target activity effectively. (**A**) A schematic view of Cas13d pooled screen experiment. The library targets both protein-coding genes and non-coding RNAs. (**B**) A schematic view of DeepCas13. It extracts spatial-temporal features of sgRNA sequence and RNA secondary structure using convolutional recurrent neural networks (**C**) Compare the predictive performance using R squared score. (**D**) A boxplot of Pearson correlation coefficient between predicted scores and LFC. The predicted scores are from 5-fold cross-validation. (**E**) A boxplot of AUC scores. Any guide with LFC < -0.5 is regarded as positive sample. (**F**) A boxplot of AUPR scores. Any guide with LFC < -0.5 is regarded as positive sample.

Combining with published Cas13d screening datasets (Supplementary Table S2), we obtained 22,599 guides that target both coding genes and non-coding elements, a unique dataset to train and evaluate predictive models for Cas13d. The large number of guides enabled us to design DeepCas13, a deep learning model to predict Cas13d sgRNA on-target efficiencies (Figure 1B). DeepCas13 uses sgRNA sequences and its predicted secondary RNA structures as input, two known features determining on-target efficiency from previous studies (*36, 37*). The output of DeepCas13 is a score, named “Deep Score” and ranging between 0 and 1, to indicate the sgRNA on-target efficiency. Features from sgRNA sequences and structures are extracted through convolutional recurrent neural networks (CRNN), which is commonly used to extract features in both spatial and temporal dimensions. These sgRNA spatial-temporal features are then concatenated in one layer, followed by a fully connected layer in neural network for prediction (see Methods for more details).

DeepCas13 is different from machine learning models developed for CRISPR-Cas9 efficiency prediction (*17, 19-21*), as it takes the predicted secondary structures as input, in addition to sgRNA sequence features. Indeed, strongly depleted sgRNAs targeting essential genes usually have higher values of minimum free energy (MFE) from secondary structures (Supplementary Figure S2A). As a result, information from the predicted secondary structures improves the precision of the model (Supplementary Figure S2B-G, *p*=0.00125).

### DeepCas13 outperforms other methods for Cas13d sgRNA efficiency prediction

We compared DeepCas13 with five conventional machine learning methods, including Random Forest (RF), XGBoost (XGB), Support Vector Machine (SVM), AdaBoost (ADA) and Gradient Boosting (GBM). For conventional machine learning methods, 185 curated features as previous described (*18*) were manually generated for training (Supplementary File 2). All methods are trained and tested from three published Cas13d screening datasets (5,726 sgRNAs in total) using five-fold cross validation.

We evaluated the performance of different models on (1) predicting the performances of all guides in the dataset, and (2) classifying guides into efficient (or non-efficient) categories. For the first evaluation (all the guides), we compared the coefficient of determination (R^2^) value and Pearson correlation coefficient (PCC) between the predicted scores and the actual log fold changes (Figure 1C, D). For the second evaluation (classification), we split all guides into two different groups (LFC <= -0.5 as positive guides and the rest as negative guides), and calculated the area under the Receiver Operator Characteristic (ROC) curve (AUC, Figure 1E) and area under the precision-recall curve (AUPR, Figure 1F) for each method. DeepCas13 has a higher R^2^ and a stronger negative PCC coefficient than other methods using 5-fold cross-validation (Figure 1C, D), indicating the better performance of DeepCas13 over other methods. Similarly, the average AUC score (from 5-fold cross-validation) of DeepCas13 was 0.87, compared with the score ranging from 0.78 to 0.85 for other methods (Figure 1E, Supplementary Figure 3). The average AUPR score, which is a better metric to evaluate the performance on unbalanced dataset (*i*.*e*., fewer positive samples), is 0.64 for DeepCas13, which is significantly higher than other approaches (ranging between 0.45 and 0.58; Figure 1F, Supplementary Figure 4), demonstrating the better performance of DeepCas13d on classifying guides into strong or weak knockdown effects.

We next compared DeepCas13 with a recently published random forest prediction model (*18*), which is denoted as RF_NBT_ below. Since RF_NBT_ only uses FACS screening datasets for training, we first evaluated whether DeepCas13, trained on FACS and additional proliferation screening datasets, outperformed RF_NBT_. For one of the two proliferation screens, denoted as A375_NBT_, we used a five-fold cross validation strategy to only use 80% of the guides (plus all guides in FACS screens) for training, and used the remaining 20% guides for performance evaluation. This process was repeated five times, and the ROC scores, Pearson correlation coefficients and precision-recall scores were recorded for both proliferations screens (Figure 2A-C and D-F for the screens in (*18*) and in this study, respectively). DeepCas13 better distinguishes efficient and inefficient sgRNAs than RF_NBT_ (Figure 2A), demonstrated by its higher AUC scores (0.84 and 0.74 for DeepCas13 and RF_NBT_, respectively). In addition, there was a much stronger correction between the predicted Deep Scores and LFC distribution (Figure 2B, PCC=-0.46 and -0.28 for DeepCas13 and RF_NBT_, respectively). For the precision and recall values (Figure 2C), most of the sgRNAs with high predicted scores from DeepCas13 were indeed efficient (*i*.*e*., high precision), and the average AUPR score for DeepCas13 is 0.5049, in comparison with 0.2774 for RF_NBT_ (Figure 2C). Similar trends were found when another proliferation screening dataset (screening in Figure 1A; denoted as A375_DeepCas13_) was used for training and evaluation (Figure 2D-F). Collectively, both DeepCas13 and RF_NBT_ performed well at ranking sgRNAs, with most negative cases at one end of a scale and positive cases at the other. Compared with RF_NBT_, DeepCas13 identified fewer false positive cases, indicated by its higher precision value than RF_NBT_ with the same recall value.

**Figure 2.**
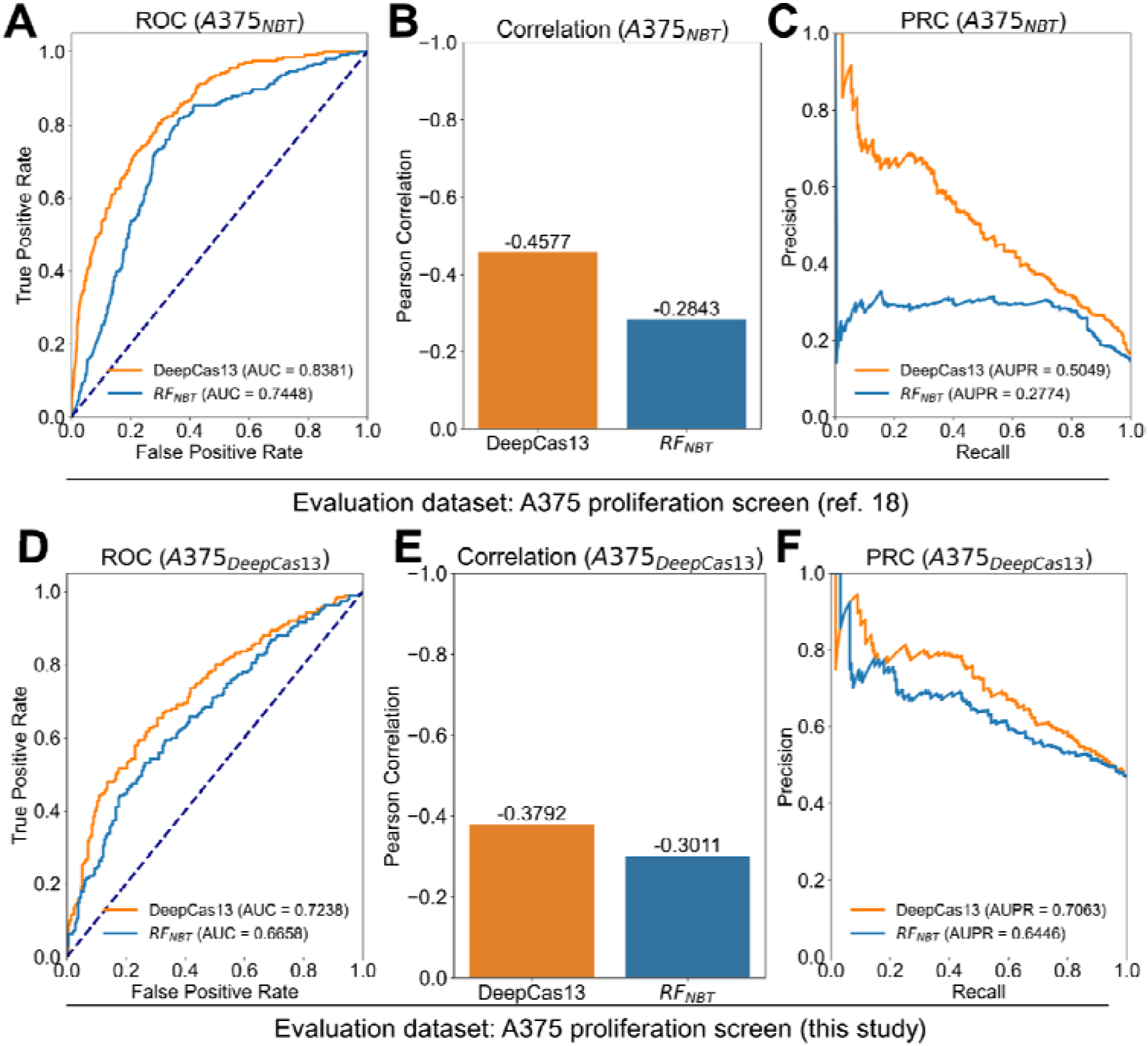
Performance comparison of DeepCas13 with state-of-the-art tool. (**A**) ROC curve comparison for the public Cas13d proliferation dataset. (**B**) Correlation comparison for the public Cas13d proliferation dataset. (**C**) PRC curve comparison for the public Cas13d proliferation dataset. (**D**) ROC curve comparison for our Cas13d proliferation dataset. (**E**) Correlation comparison for our Cas13d proliferation dataset. (**F**) PRC curve comparison for our Cas13d proliferation dataset.

We further evaluated the performance of DeepCas13 and RF_NBT_ by using one of the proliferation screening datasets as an independent validation dataset (“leave-one-dataset-out”; Supplementary Figure 5). DeepCas13 reaches higher AUROC and AUPR values than RF_NBT_ (Supplementary Figure 5), although the advantage of DeepCas13 was not as strong as in Figure 2. There was no significant difference in AUROC or AUPR values when proliferation dataset was removed from training (Supplementary Figure 5), indicating that there were few common spatial-temporal features learned between the two proliferation datasets. That may be due to the intrinsic differences within two proliferation datasets, including guide lengths (*i*.*e*., 22 bp in our study *vs* 27 bp in previous study (*18*)), whose effect may not be captured in the training dataset. Despite that, DeepCas13 consistently outperforms RF_NBT_, possibly due to the deep learning framework as well as the additional guides for training from the proliferation screens (1,398 guides for 35 essential genes in (*18*), and 3,155 sgRNAs for 94 essential genes in our study, respectively).

### DeepCas13 predicts the efficiencies of sgRNAs targeting non-coding RNAs

Having demonstrated the performance of DeepCas13 on protein-coding genes, we next evaluated whether DeepCas13 can predict the efficiency of guides targeting non-coding RNAs, including circular RNAs (circRNAs) and long non-coding RNAs (lncRNAs). Circular RNA (circRNA) is a single-stranded non-coding RNA that forms a covalently closed, continuous loop from non-canonical splicing event (*38*). circRNAs have been implicated in human physiology and diseases, although their functions are largely unclear due to a lack of adequate methods to study them (*39*). Recently, CRISPR-Cas13d has been successfully applied to study the functions of circRNAs in a screening manner (*40, 41*), providing a novel yet efficient approach to systematically investigate circRNA functions.

We first applied DeepCas13 to one circRNA screening dataset (*40*) where >□2500 human hepatocellular carcinoma-related circRNAs were screened. DeepCas13 successfully distinguished efficient sgRNAs from inefficient sgRNAs (Figure 3A, AUC score is 0.7492), although the area under the precision-recall curve (AUPR) was quite low for all the guides (Figure 3B). Since most of the circRNAs are considered to be nonfunctional, the low AUPR score may be because of the majority of the guides that have little (or no) change in abundance, even if they are predicted as efficient. Therefore, we only focused on guides that target top negative selected circRNAs (identified from the MAGeCK algorithm), and found the AUPR score greatly increased, up to 0.61 for the top 10% negatively selected circRNAs (Figure 3B).

**Figure 3.**
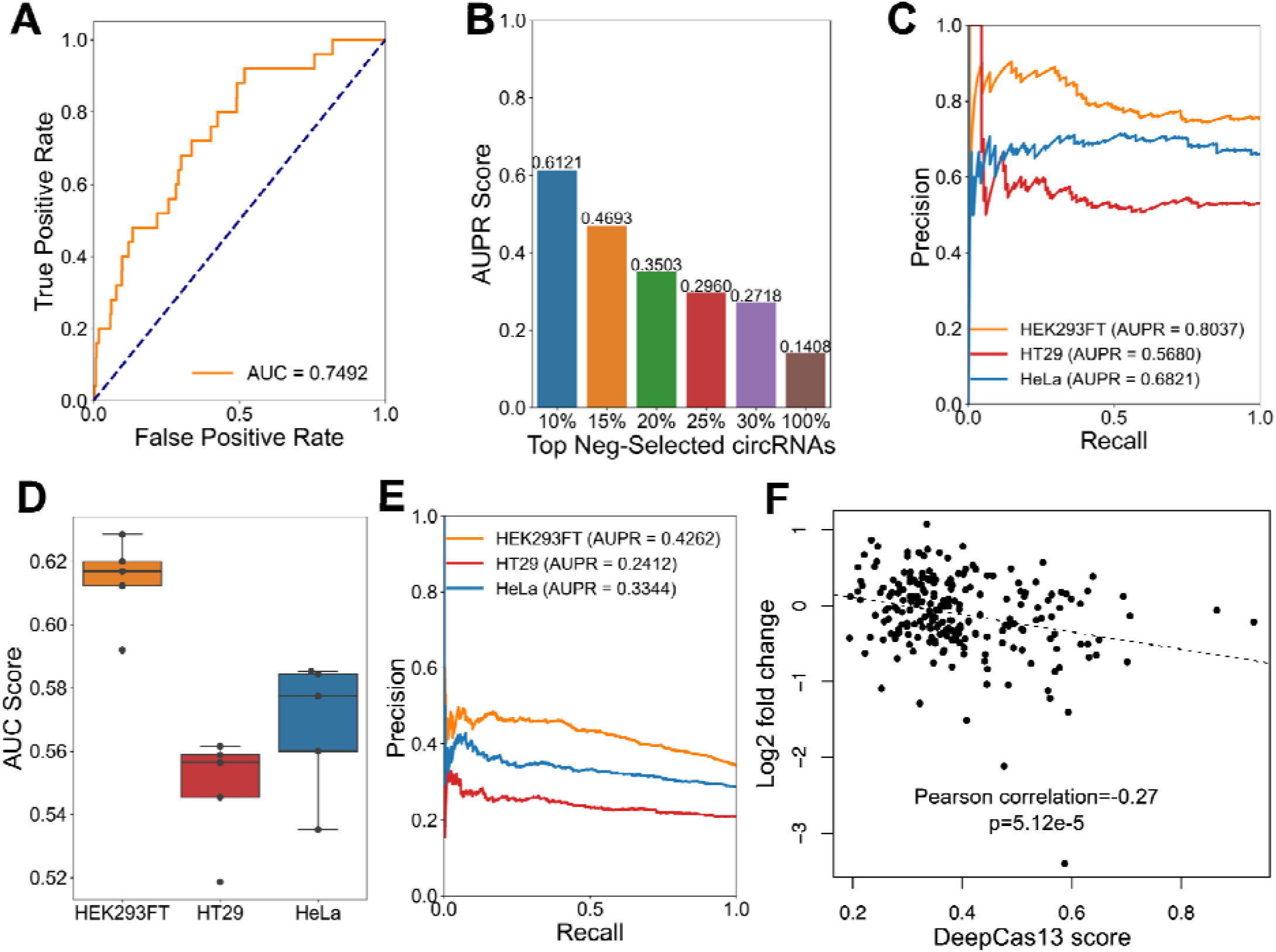
Predict activity of sgRNAs targeting non-coding sgRNAs. (**A**) ROC curve for circRNA screening dataset and the AUC score is shown in the legend. (**B**) Average precision score of the sgRNAs associated to top 10%, 15%, 20%, 25%, 30% and 100% negative selected genes. (**C**) PRC curves of the sgRNAs associated to top 50 negative selected genes for HEK293FT, HT29 and Hela cells. (**D**) A boxplot of AUC scores from 5-fold cross-validation for HEK293FT, HT29 and Hela cells. (**E**) PRC curves for HEK293FT, HT29 and Hela cells. The AUPR scores are shown in the legend. (F) The correlation between DeepCas13 predicted scores and the actual log2 fold changes for guides targeting top 20 negatively selected lncRNAs.

We also applied DeepCas13 to another circRNA screening study (*41*), where 3,800 sgRNAs were designed to target sequences across back-splicing junction (BSJ) sites of highly expressed human circRNAs in three different cell lines separately. Similarly, we limited our prediction to guides that target top 50 negatively selected circRNAs, and classified guides whose log fold change smaller than -0.5 as efficient guides. Overall, DeepCas13 predicts these circRNAs with high precision and recall (Figure 3C, AUPR scores are 0.8037, 0.5680 and 0.6821 for HEK293FT, HT29 and Hela cells, respectively). On the other hand, the AUC scores of all guides in different cell lines vary (Figure 3D), ranging from 0.63 (HEK293FT cells) to 0.54 (HT29 cells), possibly indicating some cell-type-specific effect that cannot be captured using sequence and RNA secondary structures. As a result, for those sgRNAs with high Deep Scores, many are actually not depleted (false positives), possibly because the corresponding circRNAs are not functional (Figure 3E, AUPR scores are 0.4262, 0.2412 and 0.3344 for HEK293FT, HT29 and Hela cells, respectively).

Next, we applied DeepCas13 to guides targeting 234 lncRNAs in our Cas13d screen (Figure 1A). Since not all the lncRNAs affect cell viability, we identified top 20 negatively selected lncRNAs using the MAGeCK algorithm (*35, 42*), and examined their predicted Deep Scores and the log fold change values (Figure 3F). The correlation between DeepCas13 scores and the actual log fold changes is strong (Pearson correlation coefficient=-0.27, *p* value=5.12e-5), demonstrating a good prediction power for DeepCas13 on lncRNAs. Overall, DeepCas13 demonstrated a satisfactory performance in predicting the on-target activity of sgRNAs targeting circRNAs and lncRNAs.

### Studying the off-target viability effect using machine learning approaches

We next investigated the off-target viability effect of Cas13d, or the unintended effect of Cas13d on cell viability, by examining guides that target known non-essential genes (derived from RNAi or CRISPR/Cas9 screens (*33, 34*) and are confirmed as non-essential in A375; Figure 4A; Supplementary Figure 1A). The rationale is that, since targeting these genes are confirmed to have little (or no) effect on cell proliferation and viability, the strong depletion of guides targeting these genes should come from its effect on cell viability, possibly by cleaving unintended RNA molecules (off-target RNAs or collateral RNAs). Such non-specific toxicity has been reported in other genome editing tools including shRNA (*27*) and Cas9 nuclease (*28-30*), and can be systematically evaluated in a screen manner, by examining thousands of Cas9 sgRNAs targeting the non-functional elements in the genome (*31, 32*).

**Figure 4.**
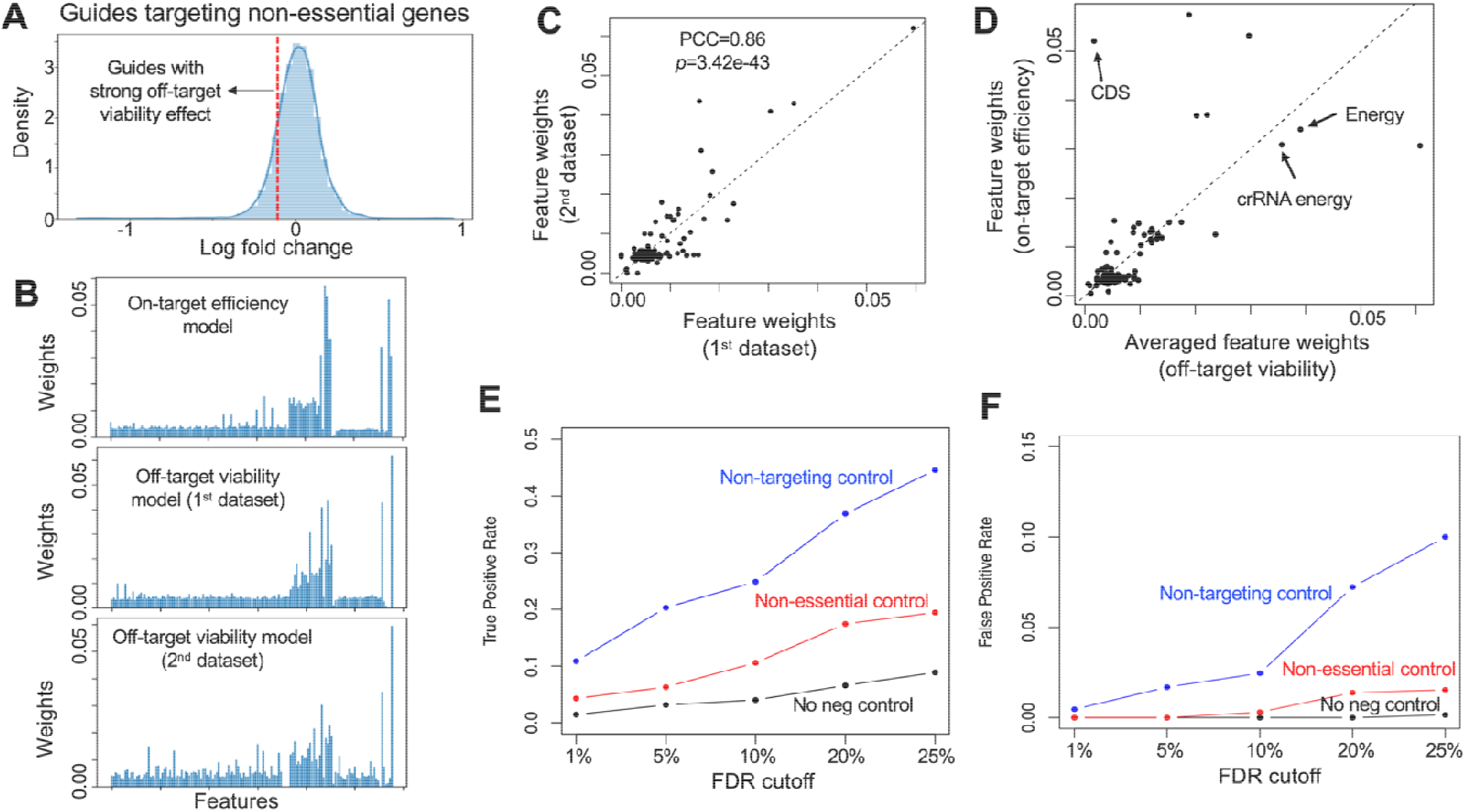
The off-target viability effect of Cas13d guides. (**A**) Study design. Guides that have strong depletion in targeting known non-essential genes are compared with other guides. (**B**) The weights of features trained by random forest by comparing guides with strong vs. weak depletion in essential genes (“on-target model”, top) and non-essential genes (“off-target viability model”, middle and bottom. (**C**) The Pearson correlation of feature weights across two different datasets. (**D**) Comparing feature weights across on-target and off-target model. (**E**) The true positive rate (in identifying known essential genes) with different FDR cutoff using different controls: no controls, non-targeting controls or non-essential controls. (**F**) The false positive rate (in identifying non-essential genes as significant) as in (E). In (E-F), the A375 cell proliferation screens from (*18*) was used.

We identified guides that have the strongest depletion in the screen (lowest LFC value), and identified features that mostly correlate with these guides. In total, 2,893 guides targeting non-essential genes from two distinct proliferation datasets (one from published viability screen (*18*) and the other from our own Cas13d screen in Figure 1A) are examined. We used random forest (RF) to train the model and identify feature importance, as the number of training samples is too few for deep learning frameworks. Interestingly, feature weights derived from these non-essential targeting guides closely resemble features determining gRNA on-target activity, which are derived from comparing guides with strong vs. weak dropouts in essential genes using random forest (Figure 4B). These feature weights are highly consistent between the two datasets used (Figure 4C), demonstrating that the off-target viability effect of Cas13d is closely related to the guide’s on-target cleavage effect, which has been extensively studied in previous sections. Indeed, the lowest predicted energy from guide, or guides plus their direct repeat (DR) sequence (“Energy” and “crRNA energy” in Figure 4D, respectively), two known features for predicting crRNA on-target efficiency, are ranked top in both on-target and off-target activity models (Figure 4D). On the other hand, whether the guides are located on the coding region (“CDS” in Figure 4D) affects its on-target efficiency, but not off-target viability effect. In addition, in three published FACS-sorting screens that do not use viability as readout, guides that have strong on-target gene depletion (“efficient guides”) also have stronger off-target viability score (trained from non-essential genes; Supplementary Figure 6A), a demonstration that the off-target effect of guides also closely correlated with their on-target activities in other screening types.

### Controlling the off-target viability effect in Cas13d screens targeting genes and lncRNAs

The unintended off-target viability effect of Cas13d guides is similar with Cas9 guides, where its DNA cleavage activity activates DNA damage response in the host cells and affects cell viability (*30*). Such effect is also observed in the screening settings, where guides targeting amplified regions of the cells have stronger depletion effect and generates false positives in the screen (*28, 29, 43*). For this reason, Cas9 guides targeting non-essential genes or non-coding regions are recommended as a better negative control (*31-33*). To compare the choice of negative control guides in Cas13d screens (non-essential genes vs non-targeting guides vs no negative control guides used), we examined the true positive rate (in identifying known essential genes as significant) and false positive rate (in identifying known non-essential genes as significant), respectively, using different FDR cutoffs (Figure 4E-F) in one Cas13d proliferation screening dataset (*18*). We observed low false positive rate (but also low true positive rate) without using any negative controls. Similar with Cas9 screens, using non-targeting guides as controls has the highest true positive rate but also introduces a large number of false positives in the screen. In contrast, using guides targeting non-essential genes demonstrated reasonable true positive rate (Figure 4E) and also a much lower false positive rate compared with non-targeting controls (Figure 4F). Similar trends were also found on another proliferation screening dataset (Supplementary Figure 6B-C). These results demonstrated that, guides targeting non-essential genes should be used instead of non-targeting guides in Cas13d screens to reach a good balance in true positive and false positive rate.

We applied our findings to the screening data that targets both protein-coding genes and lncRNAs (Figure 5A). Comparing with default setting (not using any controls), using non-essential guides as negative controls increases the number of essential genes as well as lncRNAs that are statistically significant (Figure 5A). In A375 cells, 20 lncRNAs are identified as negatively selected with statistically significance (FDR<0.25), including those that are known to be associated with tumorigenesis (Figure 5B). For example, NEAT1 has been shown to promote the proliferation, migration and invasion of melanoma cells as well as A375 cells, by interfering the regulation of multiple microRNAs and their target genes(*44, 45*). Another example is SNHG29, whose tumorigenesis role has been reported in multiple cancer types (*46, 47*). The over-expressions of both NEAT1 and SNHG29 are all associated with poor survival in TCGA melanoma cohort (Supplementary Figure 7A, B). We further performed the same screening experiment on A549 cell line (a lung cancer cell line; Supplementary Figure 7C), and compared the lncRNA screening results with their expression levels across two different cell lines (A375 and A549; Figure 5C-D; Supplementary Figure 7D). lncRNAs that have higher expressions are more likely to be negatively selected (Figure 5C). In addition, their functions show a stronger cell type-specific effect than protein-coding genes (Figure 5D, Supplementary Figure 7E-F), consistent with previous findings that these lncRNAs may likely work in a cell type-specific manner than protein-coding genes (*48*).

**Figure 5.**
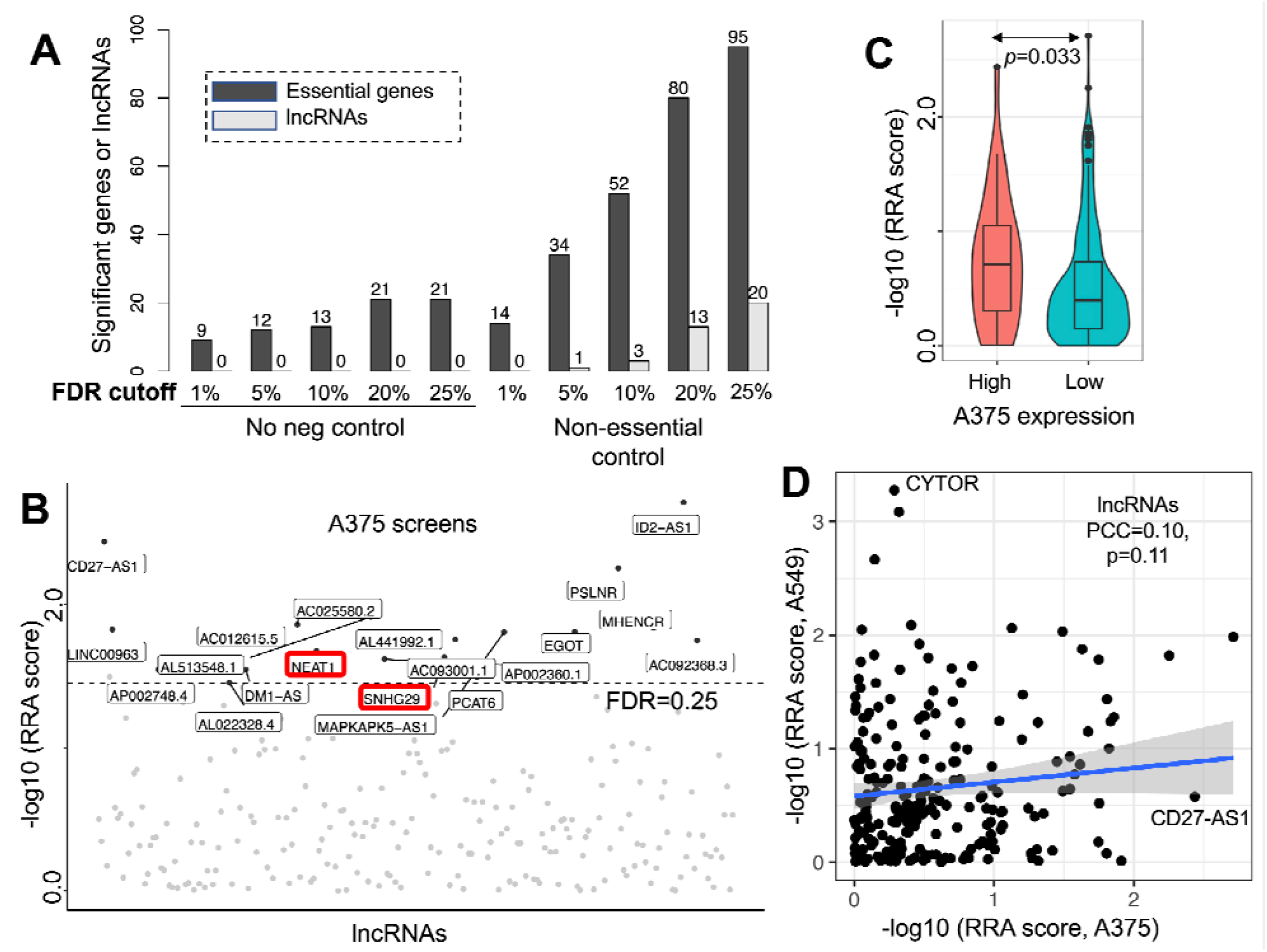
Screens for genes and lncRNAs. (**A**) The number of statistically significant genes and lncRNAs using different FDR cutoff, using either no negative control guides or using guides targeting non-essential genes (“non-essential control”) as negative controls. (**B**) Top negatively selected lncRNAs, identified from the MAGeCK algorithm. (**C**) The distribution of RRA scores, measured in the screen, of lncRNAs with high (or low) expressions in A375 cells. (**D**) The RRA scores of lncRNAs across two different cell lines. Two lncRNAs that showed distinct phenotype in two cell lines (CD27-AS1 and CYTOR) are marked.

## Discussion

A better understanding (and prediction) of the efficiency and specificity of Cas13 will accelerate the application of this new RNA editing tool to many areas. In this study, we performed CRISPR-Cas13d (CasRx) viability screens that target both protein-coding genes and non-coding RNAs. Based on the screening data (and published datasets), we designed DeepCas13, a deep learning-based model for predicting CRISPR-Cas13d on-target activity. DeepCas13 uses convolutional neural network and recurrent neural network to learn the spatial-temporal features of sgRNA sequence and RNA secondary structure. Compared with five conventional machine learning methods and a recently published state-of-the-art tool, DeepCas13 demonstrated a better performance in Cas13d sgRNA efficiency prediction (Figure 1-2). In addition, DeepCas13 performs well on both protein-coding genes and non-coding RNAs (including circular RNAs and long non-coding RNAs; Figure 3).

We also applied machine learning method (random forest) to better understand the off-target viability effect, by comparing guides that have strong vs. weak effect on non-essential genes. We found features identified from this analysis are consistent with features for on-target efficiency, implying such off-target viability effect is closely associated with the efficiency of a guide (Figure 4). Therefore, a proper negative control (targeting non-essential RNAs) should be used, especially in the proliferation/viability screens, to control for such off-target viability effect. Indeed, using non-essential control guides improves the true positive rate of a screen (compared with not using any control guides), while keeping false positive rate at a lower level (compared with using non-targeting controls).

Our Cas13d screen contains guides that target 234 lncRNAs. We demonstrated that our DeepCas13 model predicts the phenotype of guides targeting top negatively selected lncRNAs (Figure 3F), and increases the statistical power in identifying statistically significant lncRNAs using non-essential control guides (Figure 5A). The analysis also identified known and putative oncogenic lncRNAs, as well as lncRNAs that have cell type-specific functions between cell lines (Figure 5B-D). Collectively, our screening system and computational model provide a simple yet effective approach for the large-scale functional studies of non-coding RNAs (including lncRNAs).

Currently, despite the datasets used in this study, few Cas13d screening datasets are available. The power of deep learning framework is therefore limited as it performs best with a large number of training datasets. In the future, new Cas13d screens can be added as additional training samples to further improve the prediction performance of DeepCas13 and our off-target viability model. Another limitation is, DeepCas13 only focused on Cas13d (CasRx), one of the commonly used Cas13 nuclease. It is therefore unclear whether DeepCas13 works on other Cas13 proteins like Cas13a, Cas13b or Cas13XY. Once enough screening datasets (or validation datasets) for other proteins are available, DeepCas13 can be further extended to predict the efficiency of other Cas13 proteins beyond Cas13d.

## Methods

### Oligonucleotide library design

#### Gene selection

We choose 94 known “core-essential genes” and 10 known “non-essential” genes whose perturbation is known to have strong (or no) effect on cell proliferation or viability from published resources including our previous study (*33, 34*). In addition, 88 other protein-coding genes with various functions in cancer are added, including oncogenes (e.g., PIK3CA, PAK2, MYC) or tumor suppressor genes (e.g., RB1, CDKN1B).

#### lncRNA selection

We selected lncRNAs whose expressions are overexpressed in multiple cancer types in the TCGA cohort (BRCA and PRAD). Briefly, the expressions of lncRNAs for each patient sample are downloaded from TCGA, and lncRNAs are selected if they meet the following criteria: (1) their average expressions (in log2 FPKM) are greater than 2 for the 15% samples with the highest expression of this lncRNA; (2) their log2 fold changes (compared with normal samples) are greater than 0.2; and (3) their expressions (FPKM) are greater than 10 in the corresponding cancer cell lines. In addition, 25 literature-curated lncRNAs with known functions in multiple cancers are included (*49*).

#### sgRNA design

Ensembl gene and lncRNA annotations (version: GRCh38) are used to extract the sequences of corresponding genes or lncRNAs. For genes (or lncRNAs) with multiple transcripts, the transcripts that have the corresponding RefSeq ID are used. We first enumerate all possible guides (22bp) that span the entire transcript, then remove guides that (1) map to more than one location in the human genome and transcriptome (allowing up to 1 mismatch) or (2) contains BsmBI digestion sites (“CGTCTC” or “GAGACG”). The remaining guides are randomly selected if they hit the greatest number of transcripts (“common” guides) within the same gene/lncRNA. If all the guides that hit the greatest number of transcripts are selected, the remaining guides that hit the second greatest number of transcripts will be randomly selected, and so on. For essential and non-essential genes, 35 guides are designed per gene. For other genes or lncRNAs, 15 guides are designed per gene or lncRNA.

### CRISPR Library Synthesis and Construction

The pooled synthesized oligos (Synbio Technologies, China) were PCR amplified and then cloned into pLentiCasRxDR-puro vector (for expressing CasRx sgRNA) via BsmBI site by Gibson Assembly. The ligated Gibson Assembly mix was transformed into self-prepared electrocompetent Stable *E. coli* cells by electro-transformation to reach the efficiency with at least 100X coverage representation of each clone in the designed library. The transformed bacteria were cultured directly in liquid LB medium for 16∼20 hours at temperature 30°C to minimize the recombination events in *E. coli*. The library plasmids were then extracted with EndoFree Maxi Plasmid Kit (TIANGEN, Cat no. 4992194).

### Pooled Genome-wide CRISPR Screen

Human melanoma A375 cells, human non-small cell lung carcinoma A549 cells and HEK293FT cells were maintained in DMEM medium supplemented with 10% fetal bovine serum (FBS). Protein-coding gene and lncRNA-targeting plasmid libraries under lentiviral pLentiCasRxDR-puro backbone were firstly transfected along with pCMV8.74 and pMD2.G packaging plasmids into HEK293FT cells using Lipofectamine™ 2000 Transfection Reagent (Invitrogen, Cat no. 11668019) to generate Cas13d sgRNA-expressing lentivirus. Harvest virus-containing media at 72 hours post-transfection, and spin down the media at 1000 g for 5 min to remove the floating cells and cell debris. Carefully collect the virus supernatant, aliquot and store them at -80°C for further use. Test the virus titer and MOI (multiplicity of infection) before proceeding to the screen. For two-vector Cas13d cell growth screen, cas13d-expressing A375 or A549 cells were firstly generated by lentiviral infection of Cas13d (pLentiCas13d-NLS-blast) and then amplified to a number around 3×10^7^. Then, these cells were infected with Cas13d sgRNA-expressing lentiviral CRISPR library with MOI ∼0.3. Two days later, select the infected cells with puromycin (1 µg/mL for A375 cells and A549 cells) for three days to get rid of any non-infected cells before changing back to normal media. At least ∼300x coverage of cells were collected as Day 5 sample and stored at -80°C for later genomic DNA isolation. The rest of cells were continually cultured until Day 33 before harvesting as the end point sample. Genomic DNA from Day 5 and Day 33 samples was extracted. The regions encompassing the sgRNAs were firstly PCR-amplified for around 20-23 cycles with the following primer pair: pLentiCasRxDR_F1: 5’-AATGGACTATCATATGCTTACCGTAACTTGAAAGTATTTCG-3’; pLentiCasRxDR_R1: 5’-GGAGTTCAGACGTGTGCTCTTCCGATCTCCAGTACACGACATCACTTTCCCAGTTTAC-3’. The second round of PCR were employed to attach the illumina adaptors and index for around 10-12 cycles with the following primers: library_F: 5’-AATGATACGGCGACCACCGAGATCTACACTCTTTCCCTACACGACGCTCTTCCGATCTA TCTTGTGGAAAGGACGAAACACC-3’; Index_R: 5’-CAAGCAGAAGACGGCATACGAGATNNNNNNNNGTGACTGGAGTTCAGACGTGTGCTCT TCCGATCT -3’ (N(8) are the specific index sequences). These PCR products were gel purified and pooled for high-throughput sequencing to identify sgRNA abundance on illumina PE150 sequencing platform (Novogene, China).

### Cas13d screening data processing

The Cas13d screening sequence data in fastq format was processed by MAGeCK(*35, 42*) package. The MAGeCK count command was used to generate sgRNA raw read count table. For public dataset, we used its raw read count table directly if it’s provided in the original paper. Then, a Robust Rank Aggreation (RRA) algorithm was applied to normalize and rank the read counts. The sgRNA LFC was collected from the sgRNA summary table, and the top negative genes was selected from the gene summary table based on the ranking results.

### Normalize the LFC and define target value for prediction

A customized sigmoid function was used to normalize the LFC, and the calculation formula was as follows:

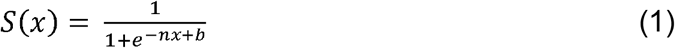

where, *x* was the LFC of a given sgRNA.

The *n* and *b* were two customized parameters to make:

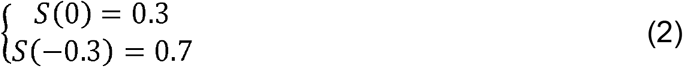

These customized parameters were based on LFC distribution of the training data, and they were designed to map effective LFC (<-0.5) closer to 1 and at the same time make the mapping range of LFC enriched region ([-0.3, 0]) as large as possible.

### Development of DeepCas13

For a given Cas13d sgRNA, we first calculate its minimum free energy (MFE) structure by ViennaRNA (*50*). The secondary structure is regarded as an equal contribution to the sgRNA sequence. In DeepCas13 model, the sgRNA structure will be a separate entry (Figure 1B). Then, the spatial-temporal features of sgRNA sequence and structure features will be extracted through convolutional recurrent neural networks (CRNN) separately. CRNN is the combination of two of the most prominent neural networks (convolutional neural network and recurrent neural network), which is designed to extract features in both spatial and temporal dimensions. It starts with a traditional 2D convolutional neural network followed by batch normalization and RELU activation. Two such convolution layers are placed in a sequential manner with their corresponding activations. The convolutional layers are then also followed by dropout (with a dropout rate of 50%) and max pooling. Next, a Long short-term memory (LSTM) layer is applied for temporal feature extraction followed by dropout (with a dropout rate of 30%) and the dense layer. Finally, the spatial-temporal features from sequence and structure are concatenated in one single layer followed by fully connected neural network for the prediction. A Deep Score for each sgRNA will be output to indicate the on-target efficiency of a specific sgRNA. The higher the Deep Score is, the more likely the sgRNA is to be effective.

### Performance comparison of DeepCas13 with conventional machine learning

Five conventional machine learning methods were used in this comparison, including Random Forest, XGBoost, Support Vector Machine, AdaBoost and Gradient Boosting. All of the methods were implemented using scikit-learn package(*51*) except XGBoost. A total of 185 features were extracted for the training based on previous study(*18*). The training data contained 5,726 sgRNAs from three Cas13d tiling screening experiments (CD46, CD55, CD71). Among them, 1,174 sgRNA were marked as positive samples as their LFC <= -0.5. We performed 5-fold cross validation to evaluate model performance on these limited training samples. To make the comparison more comprehensive, four independent indicators were used to measure the predictive performance, including R squared score, Pearson correlation coefficient, area under the ROC curve and area under the Precision-Recall curve.

### Performance comparison of DeepCas13 with start-of-the-art tool

Two Cas13d pooled proliferation screening datasets (performed on A375 cells) were used as evaluated data in this comparison separately. For each target gene, only the top 4 or bottom 4 ranked sgRNAs were left for further analysis. To increase the training set, the Cas13d tiling screening data was also combined with the training data. We also performed 5-fold cross validation for calculating Deep Scores. Both tiling screening and proliferation datasets are used for training and testing. After 5-fold cross validation, only these sgRNAs from proliferation dataset are left for further evaluation. The RF_NBT_ scores were download from cas13design Git archive directly for public A375 dataset. For A375 data from our experiment, the RF_NBT_ scores were calculated by its source code. Due to built-in filtering setting in RF_NBT_, some guides in our library can’t get a RF_NBT_ scores. So, only these sgRNAs with both Deep Score and RF_NBT_ score were left for further comparison. We compared the performance between DeepCas13 and RF_NBT_ by Pearson correlation coefficient, AUC score and AUPR score.

For leave-one-dataset-out evaluation, DeepCas13 was trained on three tiling screening datasets and one A375 pooled screening dataset, and another A375 pooled screening data was used as an independent validation dataset. We also compared the performance when trained DeepCas13 only using FACS sorting data. The AUPR score and AUC score were used to evaluate the performance.

### Performance evaluation on circRNA screening

For dataset from circRNA screening study (*40*), we performed 5-fold cross validation to get Deep Scores for these limited training sgRNAs. Any sgRNAs with LFC <= -0.5 were set as positive samples and other sgRNAs were set as negative samples. We first calculated the AUC score and AUPR score for the whole dataset. Next, the top 10%, 15%, 20%, 25% and 30% negative selected circRNAs were chosen from rank list in the gene summary table separately. Only these sgRNAs that associated to the top negative selected circRNAs were left for calculating the AUPR score.

For dataset from another circRNA screening study (*41*), top 50 negative selected circRNAs were selected from the gene summary table for HEK293FT, HT29 and Hela cell lines separately. The sgRNAs associated to these top negative selected circRNAs were used to generate Precision-Recall curve and calculate AUPR score. We also calculated the AUC scores and AUPR scores for all the sgRNAs in the library.

### Off-target viability analysis

The random forest method was implemented using scikit-learn package(*51*). All guides targeting non-essential genes in two studies are used (499 guides targeting 10 non-essential genes in our study, and 2593 guides targeting 65 non-essential genes in (*18*), respectively). We compare guides that have the strongest dropout (lowest 25% log fold change value) with the rest of the guides.

### lncRNA expression and survival analysis

The lncRNA expressions from A375 and A549 cells are from the Cancer Cell Line Encyclopedia (CCLE) and are downloaded from DepMap (version: 2019) (*52*). The survival analysis of lncRNAs is performed using “The Atlas of Noncoding RNAs in Cancer” (TANRIC) database (*53*) and DrBioRight platform(*54*).

## Acknowledgements

We would like to thank all members of the Li and Fei lab for the valuable discussion of the results.

## Author’s contributions

WL and TF conceived and designed the study. WL, XC and ZL designed the library. ZL and TF performed Cas13d screens. XC, WL, ZL processed, analyzed and interpreted the data. XC and RS designed and implemented the DeepCas13 website with the help of JP. XC, WL, and TF wrote the manuscript with the help of other authors. WL and TF supervised the study. All authors read and approved the final manuscript.

## Availability of Data and Materials

The Cas13d screening data is deposited into Gene Expression Omnibus (GEO) under the accession number GSE183256. The DeepCas13 algorithm can be accessed at http://deepcas13.weililab.org.

## Ethics Approvals

Not applicable.

## FIGURE LEGENDS

**Figure S1.**
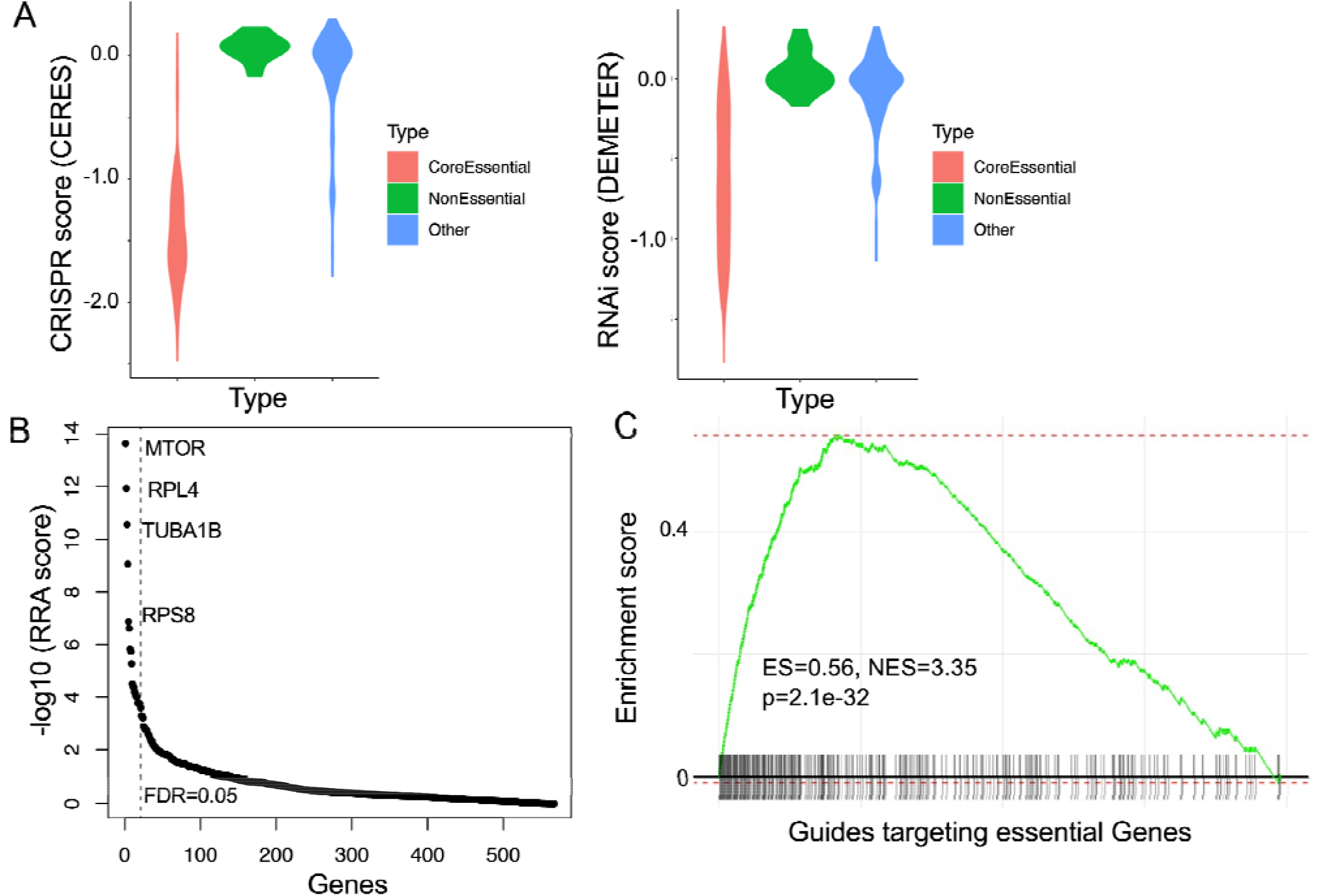
Summary of CRISPR-Cas13d screens. **(A)** The scores of essential genes, non-essential genes and other genes in A375 RNAi screens and CRISPR screens in DepMap. CRISPR scores are calculated using CERES, while RNAi scores are calculated using DEMETER in DepMap. (**B**) The RRA score distribution of negative selection (day 33 vs day 5), reported by the MAGeCK algorithm. (**C**) The overall enrichment of guides targeting essential genes in the ranking of all the guides, estimated by Gene Set Enrichment Analysis (GSEA). ES: enrichment score; NES: normalized enrichment score.

**Figure S2.**
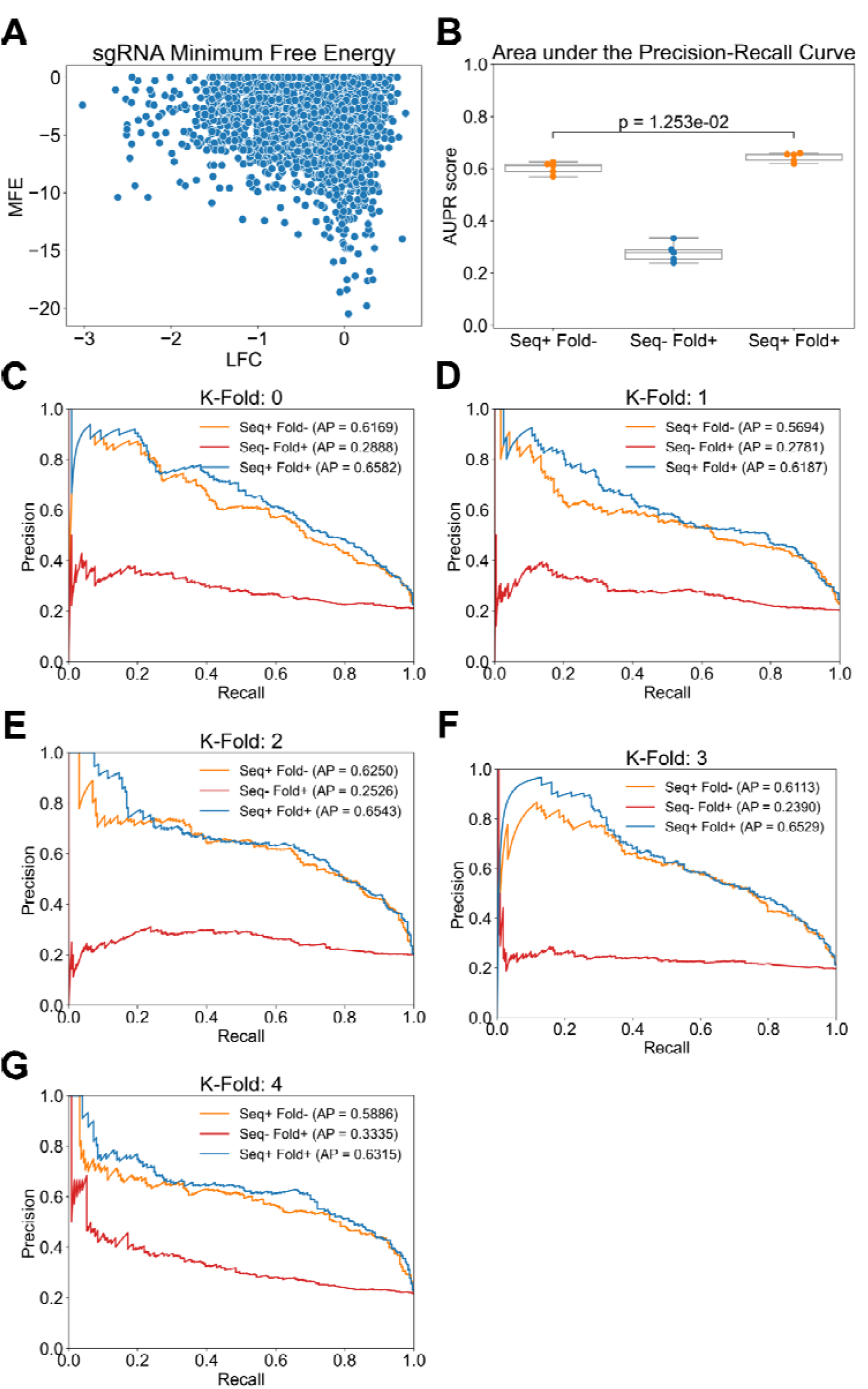
sgRNA secondary structure is beneficial to model precision. (**A**) Scatterplot shows the relationship between MFE and LFC. (**B**) Average precision comparison with or without sgRNA secondary structure. (**C-G**) Precision-recall curves from 5-fold cross-validation.

**Figure S3.**
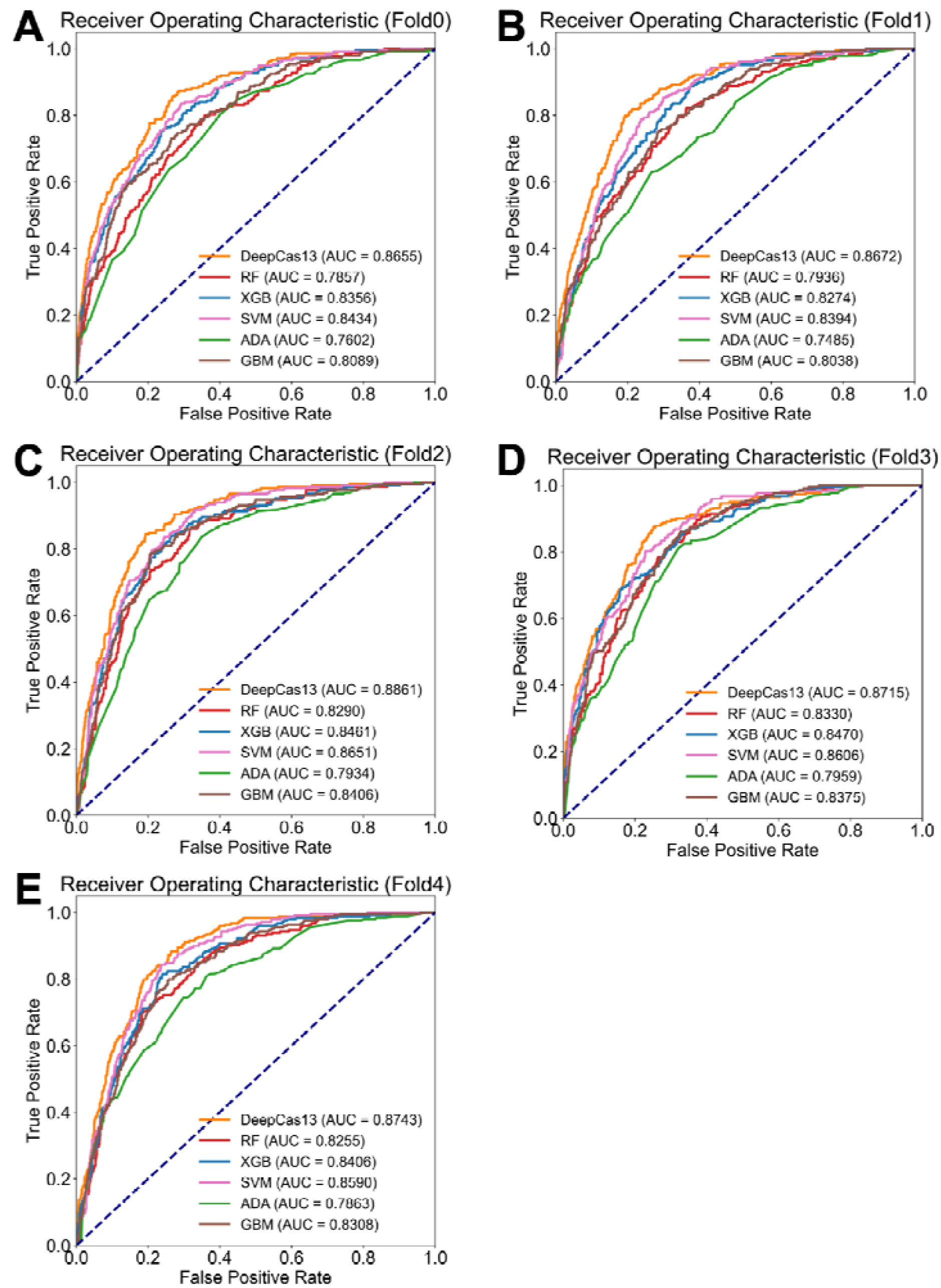
DeepCas13 can distinguish between effective and invalid sgRNAs. (**A**) ROC curves of the 1^st^ fold of validation data and the AUC scores are shown in the legend. (**B-E**) ROC curves of the 2^nd^-5^th^ fold of validation data and the AUC scores are shown in the legend.

**Figure S4.**
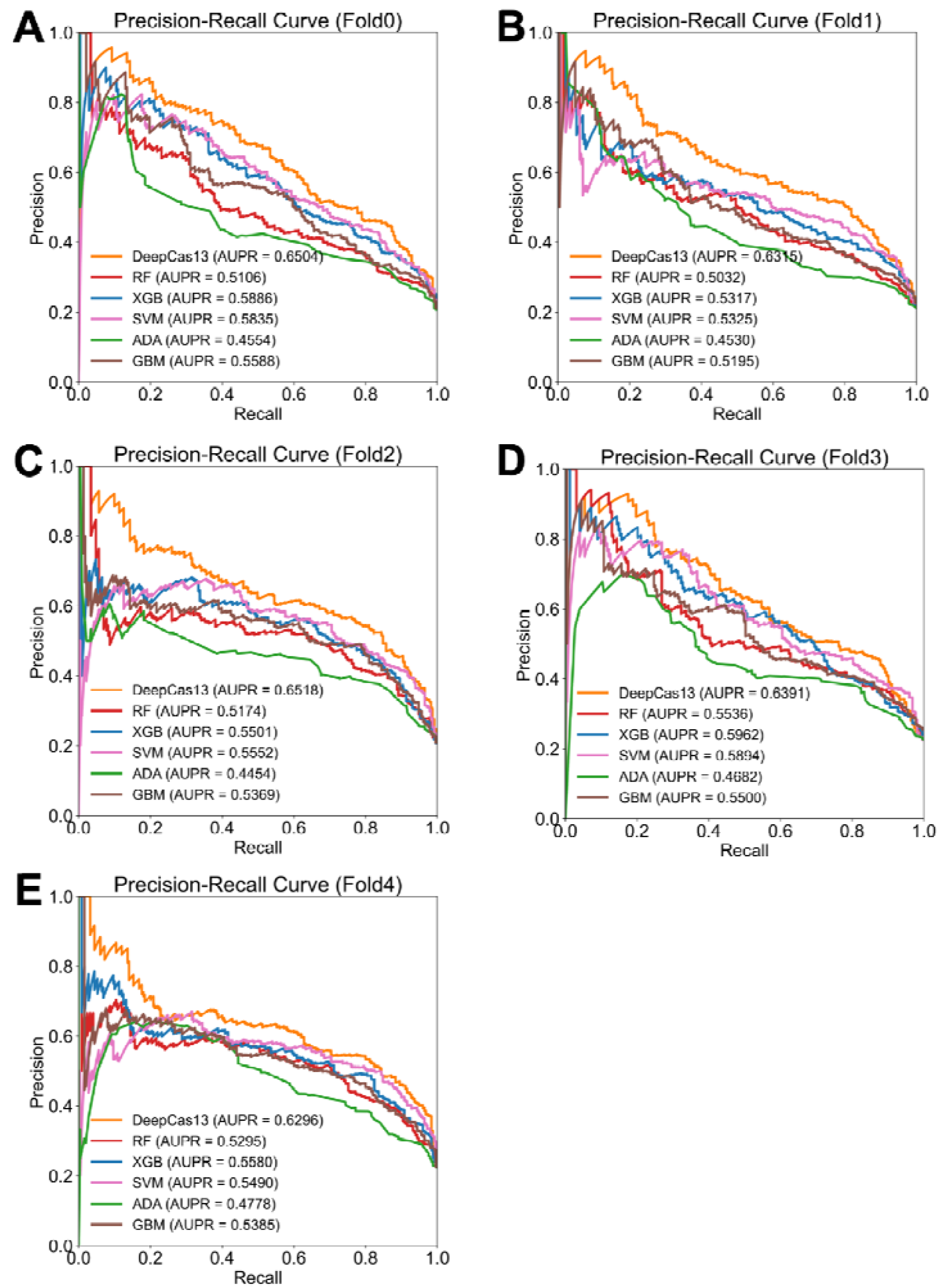
DeepCas13 can reach high prediction precision. (**A**) PRC curves of the 1^st^ fold of validation data and the AUPR scores are shown in the legend. (**B-E**) PRC curves of the 2^nd^- 5^th^ fold of validation data and the AUPR scores are shown in the legend.

**Figure S5.**
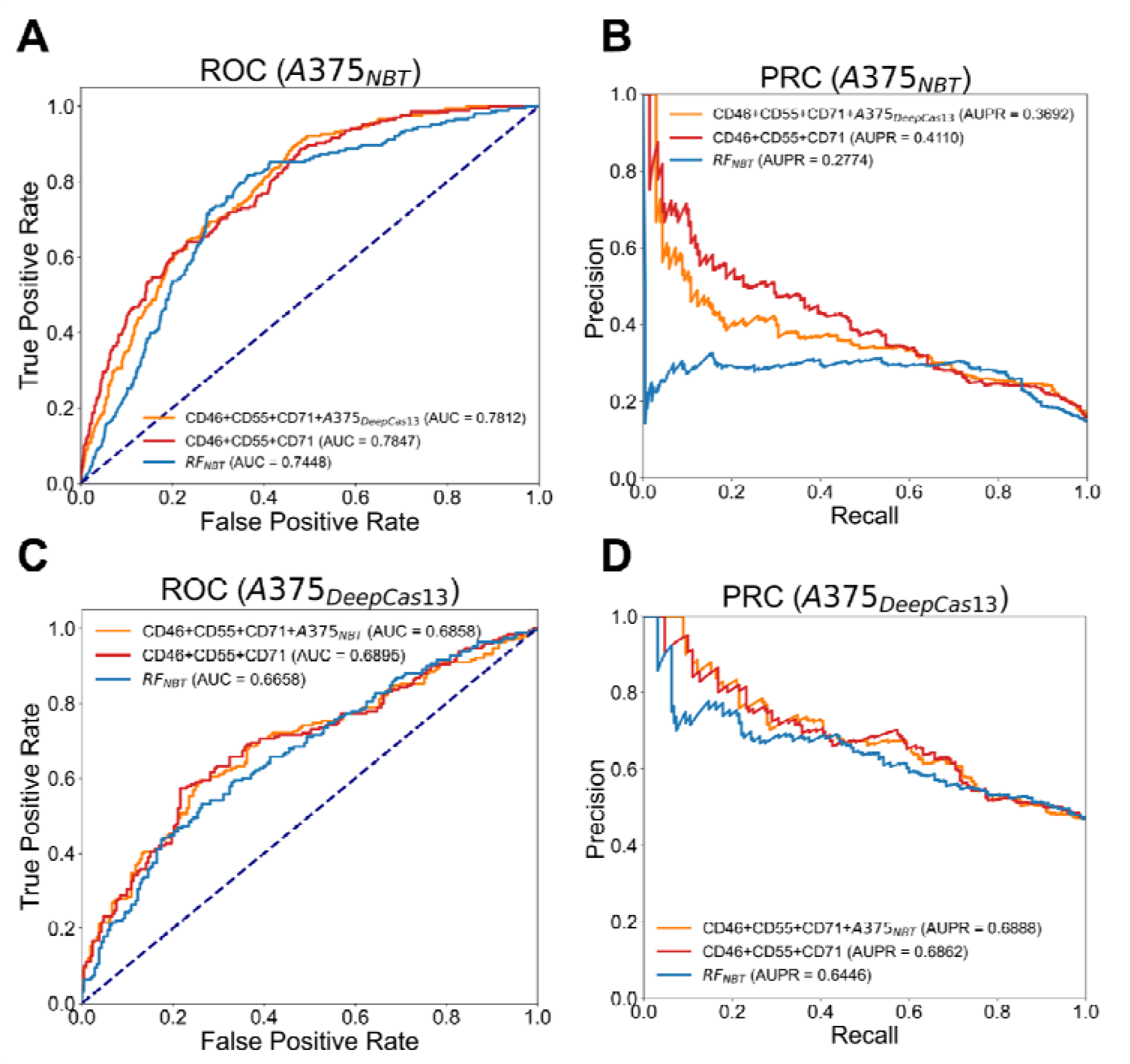
Leave-one-dataset-out evaluation. (**A**) ROC curve comparison for the public Cas13d proliferation dataset. Orange curve means both FACS sorting data and proliferation data are used for the training. Red curve means only FACS sorting data is used for the training. Blue curve show the performance of the existing tool. (**B**) PRC curve comparison for the public Cas13d proliferation dataset. (**C**) ROC curve comparison for our Cas13d proliferation dataset. (**D**) PRC curve comparison for our Cas13d proliferation dataset.

**Figure S6.**
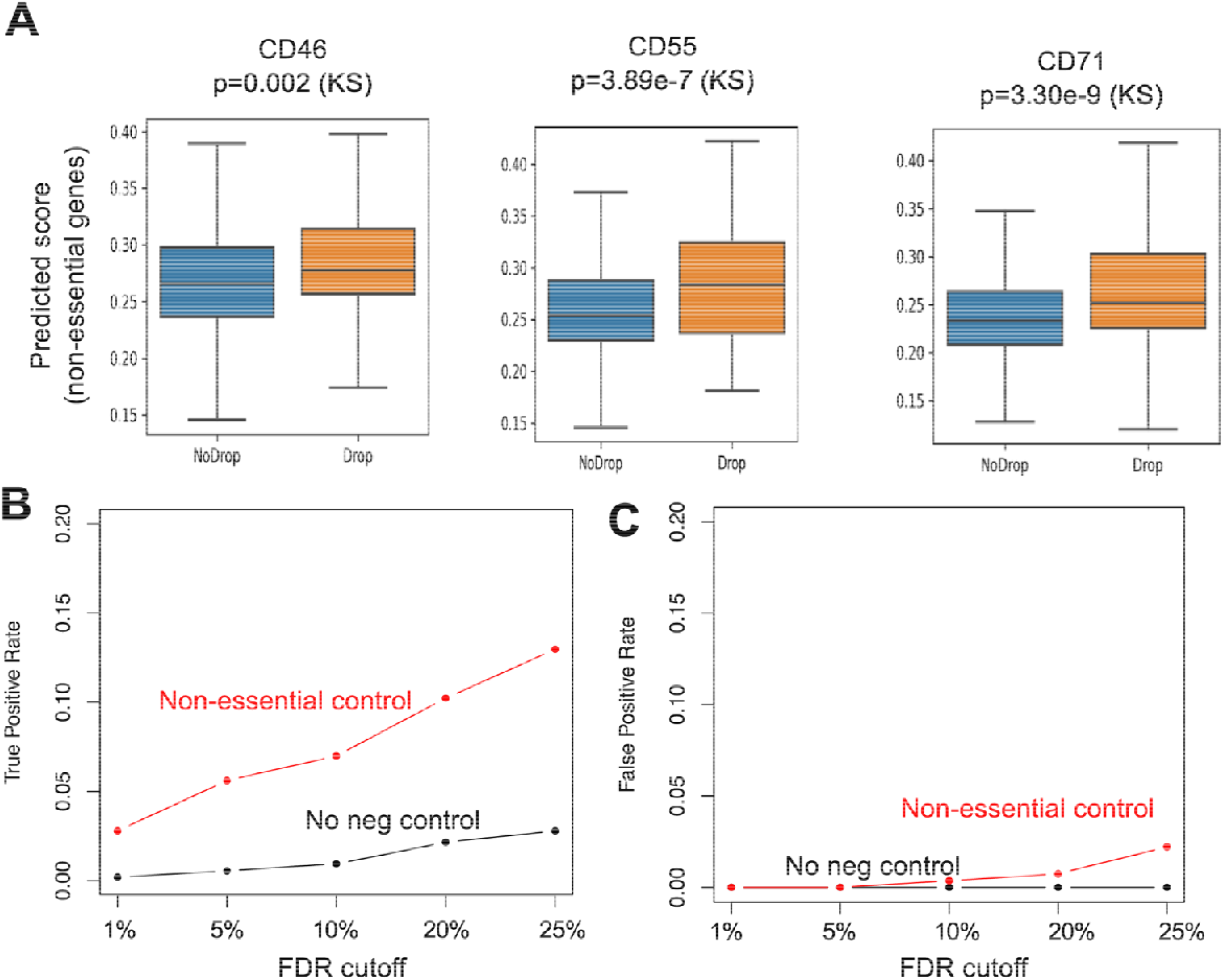
(**A**) The predicted off-target viability score (using non-essential genes) on guides that showed strong on-target knockdown (“Dropped” guides) of target genes in FACS-based Cas13d screens vs. other guides. The “Dropped” guides are guides that demonstrated strongest dropout in GFP+ population. (**B**) The true positive rate (in identifying known essential genes) with different FDR cutoff using different controls: no controls or non-essential controls. The A375 screens in our study was used. Non-targeting control analysis was not included as the library did not contain non-targeting controls. (**C**) The false positive rate (in identifying non-essential genes as significant) as in (E).

**Figure S7.**
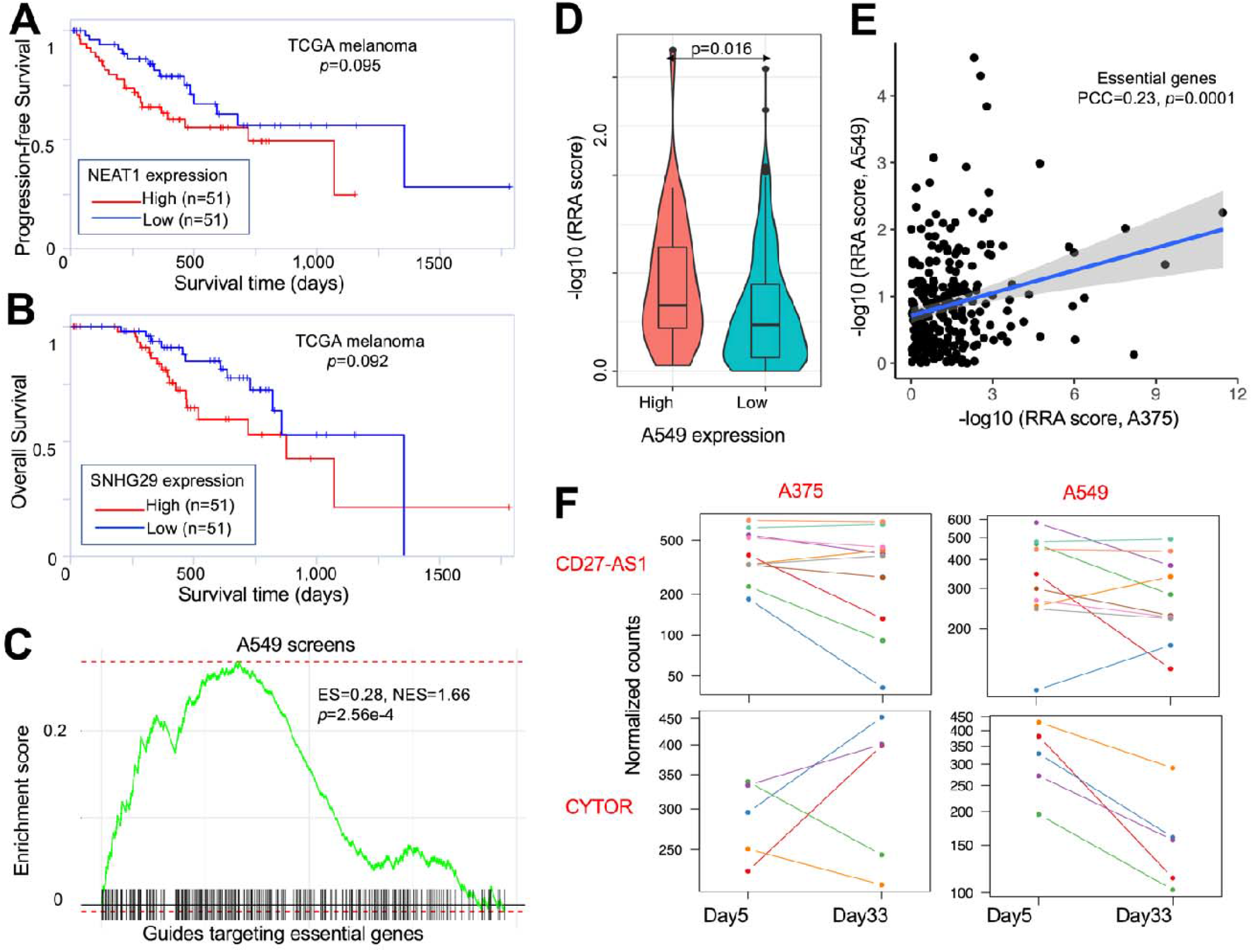
lncRNA screening analysis. (**A**) The gene set enrichment analysis (GSEA) of guides targeting essential genes, in all the guides in the screen. Guides are ranked by their negative selection in A549 cells. (**B**) The distribution of RRA scores, measured in the screen, of lncRNAs with high (or low) expressions in A549 cells. **(C-D)** The survival analysis of NEAT1 and SNHG29 expression in TCGA melanoma cohort (Skin Cutaneous Melanoma). The progression-free survival of NEAT1 and overall survival of SNHG29 is used. The analysis is performed in TANRIC platform(*53*). The *p* value is calculated using linear regression (LR). (**E**) The RRA scores of essential genes across two different cell lines. (**F**) The sgRNA changes of two lncRNAs (CD27-AS1 and CYTOR) across two different cell lines.

**TABLE S1.** The QC metrics reported by the MAGeCK algorithm

| Sample | Reads | Mapped | Percentage | Total sgRNAs | Reads per sgRNA | Zero counts | Gini Index |
| --- | --- | --- | --- | --- | --- | --- | --- |
| A375_Day5_R1 | 6,923,422 | 5,745,864 | 0.8299 | 10,829 | 530.60 | 1 | 0.04066 |
| A375_Day5_R2 | 4,540,478 | 3,772,583 | 0.8309 | 10,829 | 348.38 | 1 | 0.0451 |
| A375_Day35_R1 | 4,944,792 | 4,127,582 | 0.8347 | 10,829 | 381.16 | 2 | 0.05567 |
| A375_Day35_R2 | 6,114,984 | 5,033,558 | 0.8232 | 10,829 | 464.82 | 3 | 0.05655 |

**TABLE S2.** Summary table of training data used in this study

| Dataset | sgRNA number | Target | Source |
| --- | --- | --- | --- |
| Cas13d A375 pooled screens data | 10,830 | essential/non-essential genes, lncRNAs | this study |
| Cas13d tiling screens data | 5,726 | CD46, CD55, CD71 | PMID: 32518401 |
| Cas13d A375 pooled screens data | 1,398 | essential genes | PMID: 32518401 |
| Cas13d circRNA screens data | 3,800 | circRNAs | PMID: 33288960 |
| Cas13d circRNA screens data | 845 | circRNAs | PMID: 33478577 |

